# Determining the microbial species content in tissue from Alzheimer’s and Parkinson’s Disease patients

**DOI:** 10.1101/2023.10.25.563977

**Authors:** Rodrigo Leitao, Iam Ut Wan, Matthew C. Fisher, Johanna Rhodes

**Affiliations:** MRC Centre for Global Infectious Disease Analysis, School of Public Health, Imperial College London, London W12 0BZ, UK; Department of Medical Microbiology, Radboudumc, Nijmegen, the Netherlands

## Abstract

The aetiology of Alzheimer’s disease (AD) and Parkinson’s Disease (PD) are unknown and tend to manifest at a later stage in life; even though these diseases have different pathogenic mechanisms, they are both characterized by neuroinflammation in the brain. Links between bacterial and viral infection and AD/PD has been suggested in several studies, however, few have attempted to establish a link between fungal infection and AD/PD. In this study we develop and describe a nanopore-based sequencing approach to characterise the presence or absence of fungi in both human brain tissue and cerebrospinal fluid (CSF). This approach detects fungal DNA in human brain and CSF samples even at low levels, whereas our quantitative polymerase chain reaction (qPCR) assay (FungiQuant) was unable to detect fungal DNA in the same samples. Comparison against kit-controls showed ubiquitous low-level fungal contamination that we observed in healthy human brains and CSF as well AD/PD brains and CSF. We use this technique to demonstrate the presence of fungal DNA in healthy human brains as well as AD/PD brains, with *Alternaria spp*., *Colletotrichum graminicola*, and *Filobasidium floriforme* as the most prominent species. In addition, antibiotic resistant *Pseudomonas spp*. was identified within the brain of an AD patient. Our method will be broadly applicable to investigating potential links between microbial infection and AD/PD.

## Introduction

Alzheimer’s disease (AD) and Parkinson’s Disease (PD) are two of the most common neurodegenerative diseases in the world. Patients with AD or PD display a loss of neurons in the central nervous system (CNS), however, the mechanism by which this occurs has yet to be fully understood^1^. AD is the leading cause of dementia, characterised by progressive cognitive impairment, memory deficits, and loss of independence^2^, with over 520,000 people in the UK being affected by dementia caused by AD^3^. Parkinsonism is a clinical syndrome characterized by slowness of movement, rigidity, tremor, and postural instability^4^. Causes of Parkinsonism may include idiopathic PD and Lewy body dementia (LBD), with PD being the most common form of Parkinsonism^4^ and the second most common type of degenerative dementia in patients aged over 65 years^5^. Prevalence of AD, PD, and dementia is increasing steadily, and identified risk factors include age, diabetes, hypertension, head injury, chronic kidney disease, depression, chronic obstructive pulmonary disease, stroke, genetic susceptibility, and environmental factors, with further risk underpinning the aetiology of these diseases remaining largely unknown^2,6–8^.

Recently, the potential role of microbial infection as an AD/PD risk factor or mediator of progression has been gaining attention due to the recognition of inflammation as a prominent feature of AD/PD, alongside associative evidence for elevated levels of proinflammatory cytokines in AD and PD patients and increased AD or PD risk conferred by variants in numerous immune-related genes^9–11^. Population-level evidence for microbial risk factors has been observed, with tuberculosis patients in Taiwan showing a 1.38-fold increased risk of developing PD compared to non-tuberculosis controls^12^. In addition, it has been hypothesised that bacteria such as *Porphyromonas gingivali*s and *Chlamydia pneumoniae* may cross the blood-brain barrier (BBB) and reach the CNS where it might cause neurological damage by inducing neuroinflammation^13^. Moreover, a study conducted by Bu et al.^14^, found that the infectious burden (IB) consisting of herpes simplex virus type 1 (HSV-1), *Borrelia burgdorferi, Chlamydophila pneumoniae* and *Helicobacter pylori* have the capability of causing cognitive decline and that such infections are linked with AD.

However, in contrast to viruses and bacteria, the role of fungi in AD/PD and dementia has received less attention. The potential involvement of fungi should not be overlooked as many species are capable of infecting the CNS and neuropathology may be mediated by subsequent inflammation^15^. More direct evidence supporting the potential role of fungal infection in AD/PD originates from a series of studies identifying fungal proteins and DNA in AD/PD patients’ brains and CSF^16–22^. Although these findings appear promising, the presence of fungal proteins and DNA in AD/PD brains and CSF is insufficient for establishing the role of infection in the cause or progression of AD/PD as it is unknown whether the fungi were viable and actively contributing to disease pathology^23^. Furthermore, fungal material was also detected in some control individuals^19–21^, which may be expected as increased BBB permeability is also associated with normal healthy ageing^24^ and there is the possibility of brain microbiomes being present^23^. For these reasons, it is necessary to develop methods to further establish the potential involvement of fungi in the CNS of AD/PD patients and to establish whether the fungi are potentially pathogenic as well as contributing to neurodegeneration via inflammatory pathways. This study aims to develop and validate methods to investigate the above hypotheses using the MinION platform from Oxford Nanopore Technologies to identify the species present.

## Methods

### Mice tissue

Mice brains (Cftrtm1Unc CF mice) were obtained from the Darius Armstrong James laboratory (Department of Infectious Diseases, Imperial College London). Brain tissue from naive mice was used to optimize the DNA/RNA extraction method, quantitative polymerase chain reaction (qPCR), polymerase chain reaction (PCR) and nanopore sequencing.

### Human tissue

Human tissue/CSF was obtained from the Imperial College Healthcare Tissue Bank (ICHTB), Parkinson’s UK Brain Bank (PB), and London Neurodegenerative Diseases Brain Bank (LNDBB) (see Table 1).

**Table 1:** Human brain tissue and CSF obtained from ICHTB (Imperial College Healthcare Tissue Bank), PB (Parkinson’s UK Brain Bank) and LNDBB (London Neurodegenerative Diseases Brain Bank). NA = not available.

| ID | Brain/CSF | Clinical diagnosis | Brain Location | Age | Brain supplied |
| --- | --- | --- | --- | --- | --- |
| C1 | Brain | Normal frozen tissue | NA | 59 | ICHTB |
| C2_1 | Brain | Normal frozen tissue | NA | 30 | ICHTB |
| C2_2 | Brain | Normal frozen tissue | NA | 30 | ICHTB |
| C2_3 | Brain | Normal frozen tissue | NA | 30 | ICHTB |
| AD1_1 | Brain | Alzheimer's disease | Cerebellum | 86 | PB |
| AD1_2 | Brain | Alzheimer's disease | Frontal cortex | 86 | PB |
| C3_1 | Brain | Control case (extensive amyloid angiopathy) | Frontal cortex | 90 | LNDBB |
| C3_2 | CSF | Control case (extensive amyloid angiopathy) | CSF | 90 | LNDBB |
| AD2_1 | Brain | Alzheimer's disease | Frontal cortex | 87 | LNDBB |
| AD2_2 | CSF | Alzheimer's disease | CSF | 87 | LNDBB |
| PD1 | Brain | Parkinson's disease | Frontal cortex |  | PB |
| AD3 | Brain | Alzheimer's disease | Frontal cortex | 85 | PB |
| C4 | Brain | Control case (ovarian cancer) | Frontal cortex | 61 | PB |
| PD2 | Brain | Parkinson's disease | Frontal cortex | 61 | PB |
| AD4 | Brain | Alzheimer's disease | Frontal cortex | 92 | LNDBB |

### DNA extraction

Five different DNA extraction methods were tested in parallel (supplementary file S1). Amongst these methods, the DNeasy PowerSoil Pro Kit was selected as it provided better performance. DNA was extracted from 25 mg brain samples using the DNeasy PowerSoil Pro Kits (Qiagen, Germany, 47014) following the manufacturer’s protocol with the following modifications: a program with 6 repeats of bead beating at 4.5 m/sec was performed for 45 seconds using the FastPrep-24™ 5G (Thermo Fisher). Samples were incubated for two minutes on ice after the third repeat. The solution C6 was replaced with elution buffer AE (19077) and the elution volume was decreased to 36 μL with an incubation step of 10 minutes. Nuclease-free water and PBS buffer were extracted with the samples to control for potential reagent contamination.

### Handling brain/CSF samples in the laboratory

To avoid sample contamination, the handling of human brain and CSF samples as well as DNA/RNA extractions were performed in a safety cabinet (class 2) equipped with a HEPA filter. The safety cabinet, pipettes, tube racks, and other lab consumables were subjected to a treatment of UV light for 30 min before the handling of the samples. In addition, all pipettes and working surfaces were decontaminated with DNA AWAY decontaminant and 10% ChemGene. Disposable sterile blades were used to perform the slicing of the human brains and sterile filtered tips were used to handle the CSF samples. All Eppendorf, PCR tubes, and beads (bead beating) used in this study were DNA/RNA free to avoid the introduction of contaminants. All DNA and RNA extractions had negative controls (nuclease free-water and PBS) and kit controls.

### Cerebrospinal fluid (CSF)

CSF was extracted by using the DNeasy Blood & Tissue Kit (Qiagen, 69504) with the following modifications: 500 μL of CSF was centrifuged for 10 min at 13,000 rpm and the supernatant (300 μL) was discarded. CSF was then resuspended with remaining fluid (200 μL). The volume of buffer AE was reduced to 36 μL.

### RNA extraction

RNA was extracted from 25 mg brain samples using the ZymoBIOMICS™ RNA Miniprep Kit (Zymo Research R2001) following the manufacturer’s protocol with the following modifications: 3 repeats of bead beating at 4.5 m/sec was performed for 45 seconds using the FastPrep-24™ 5G (Thermo Fisher) and the final elution volume was decreased to 40 μL.

### Reverse transcription qPCR

RNA extracted from AD/PD and control patients were reverse transcribed into cDNA using the High-Capacity cDNA Reverse Transcription Kit (Thermo Fisher Scientific, 4368814). Followed by qPCR (for more information see qPCR assay section).

### Determining limit of detection (LOD) for qPCR

*Aspergillus fumigatus, Candida auris*, and *Cryptococcus* spp. were cultured in Sabouraud Dextrose Agar (SDA) containing chloramphenicol at concentration of 100 mg/L. *A. fumigatus* was incubated for 3 days at 45°C and *C. auris* and *Cryptococcus. spp*. were incubated for 5 days at 37°C. After the incubation period, *A. fumigatus, C. auris*, and *Cryptococcus spp*. were sub-cultured into T25 Nunc EasYFlasks containing SDA. The harvesting of fungal spores for the above species was performed by adding 10 ml of sterile 0.1% Tween into the T25 flasks, followed by vigorous agitation and finally filtration through a 10 ml sterile syringe filled with glass wool. Spore counting was performed using a Hemocytometer Counting Chamber (Neubauer-improved bright line Marienfeld, Germany). The spore suspensions of each fungal species were diluted to 2 million spores/ml. A combined 1 million spore (absolute number) mock community containing equal proportions of *A. fumigatus, C. auris* and *Cryptococcus spp*. was made. To mimic the presence of fungi in infected brains, 25 mg mice brain samples were spiked with 1 ml of 10-fold serial dilutions of the mock community from 10,000 spores to 1 spore (absolute number). Spiked brains were vortexed for 15 seconds and centrifuged at 13,000s rpm for 5 minutes. Supernatant was removed and mice brains were subjected to DNA purification procedures using the DNeasy PowerSoil Pro Kits (Qiagen, Germany, 47014). DNA purification was followed by qPCR as described below. The lowest spore concentration detected by qPCR was considered the LOD.

### Quantitative polymerase chain reaction (qPCR) assays (FungiQuant)

To detect fungal DNA and cDNA in extracted mice and human brain and CSF samples, qPCR was performed using probes and primers (Table 2) targeting a highly conserved region of the fungal 18S ribosomal DNA (rDNA). Sequences were obtained from Liu *et al*.^25^, and FungiQuant qPCR settings are shown in Table 2.

**Table 2:**
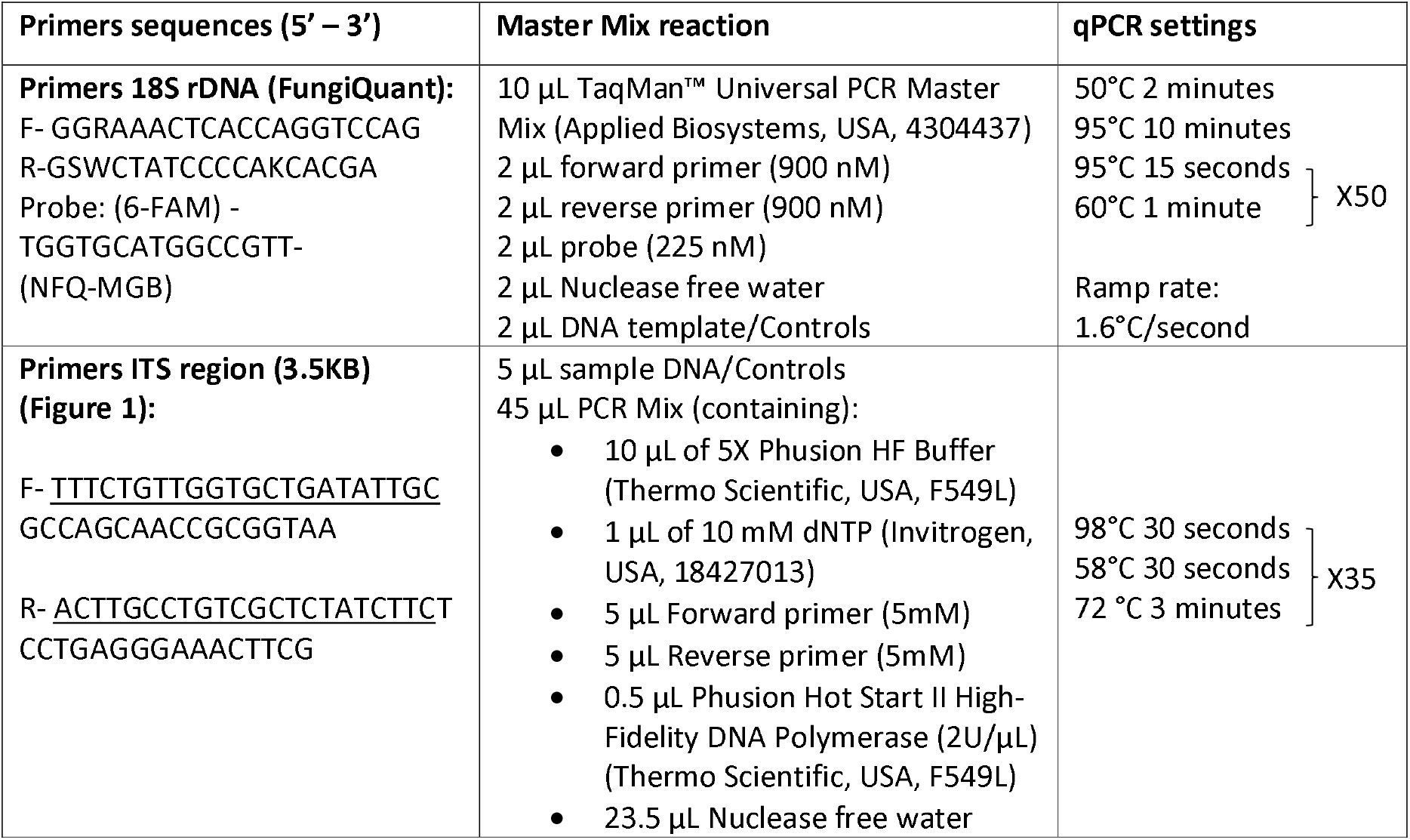
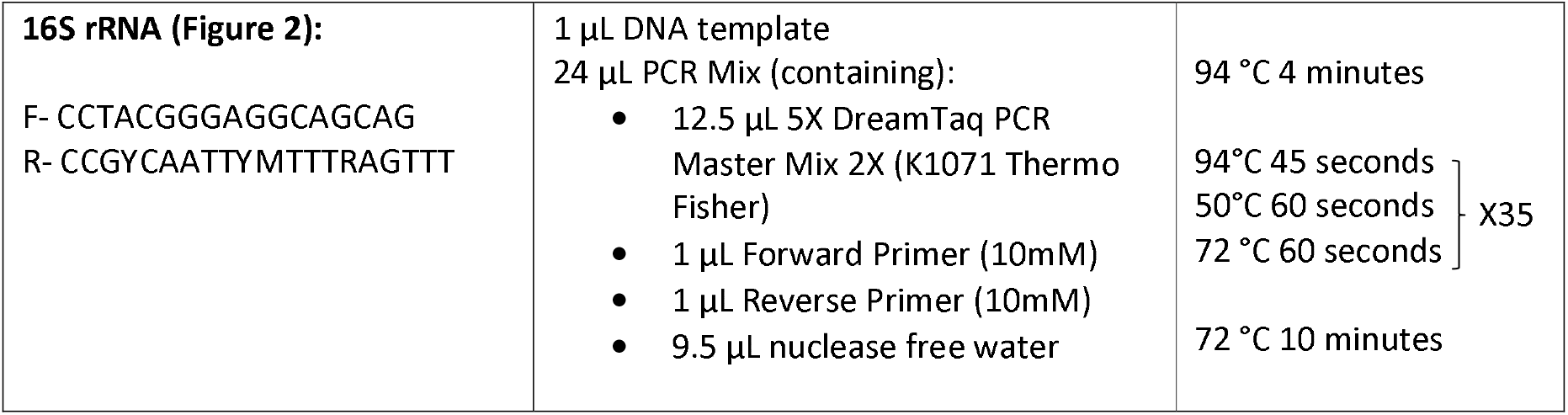
PCR/qPCR primers used in this study and its respective settings.

A standard curve was generated for each 96-well plate using 10-fold serial dilutions of Mycobiome Genomic DNA Mix (ATCC, USA, MSA-1010), containing equal proportions of *A. fumigatus, Cryptococcus neoformans, Trichophyton interdigitale, Penicillium chrysogenum, Fusarium keratoplasticum, C. albicans, Candida glabrata, Malassezia globosa, Saccharomyces cerevisiae*, and *Cutaneotrichosporon dermatis*, ranging from one million to one genome copy equivalence. qPCR on each sample was conducted in triplicate with additional no template controls (all reaction components excluding DNA) and positive controls (*A. fumigatus, C. auris* and *Cryptococcus spp*). Data was analysed using the Design and Analysis Software v2.6.0 (Applied Biosystems). In the data analysis, default thresholds were used and the average quantification cycle (Cq) of nuclease-free water, PBS, or the no template control, whichever amplifies earlier, was used as a cut-off to reduce false positive results generated by reagent contamination. Samples were defined as positive for fungi when ≥ 2/3 triplicates demonstrate amplification^25,26^ with Cq values lower than the average Cq of nuclease-free water, PBS, or the no template control.

### MinION sequencing

#### Polymerase chain reactions (PCR)

Prior to sequencing, initial PCR was performed on DNA extracted human brain and CSF to amplify fungal targets and attach sequencing barcodes. Primers targeting the fungal ribosomal operon from the V3 region of 18S rDNA to D3 region of 28S rDNA (Figure 1 and Table 2) were used. Sequences were obtained from D’Andreano & Francino^27^ and PCR settings are shown in Table 2. Negative control templates (nuclease free water) were included in all PCR runs.

**Figure 1:**
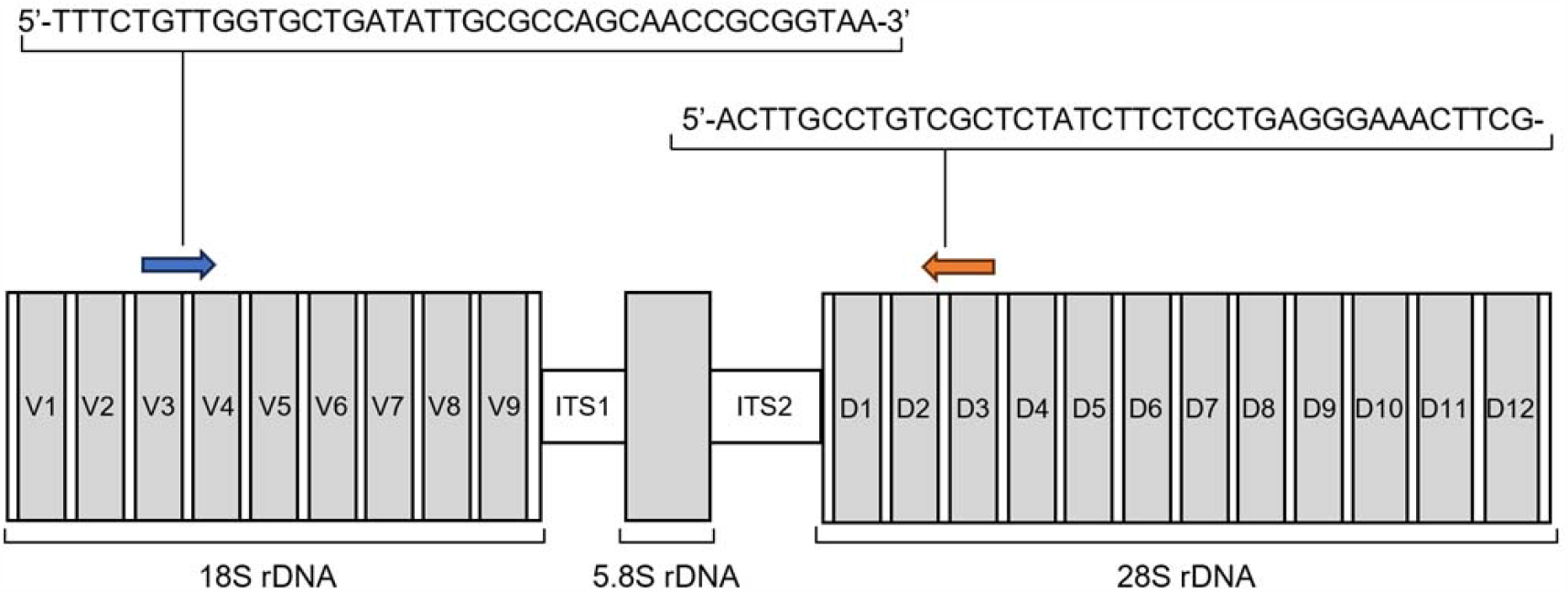
Schematic representation of the fungal ribosomal operon region to be sequenced. Forward (blue arrow) and reverse (orange arrow) primers targeting the fungal ribosomal operon and their sequences are shown. Oxford Nanopore universal tag sequences preceding the 18S and 28S rDNA-specific primer sequences are underlined. The resulting ∼3.5 kb amplicon covers hypervariable internal transcribed spacer regions 1 and 2 (ITS1 and ITS2), as well as variable domains V3-V9 of the 18S rDNA gene and D1-D3 of the 28S rDNA gene, which can refine taxonomic assignment. Figure modified from D’Andreano & Francino^27^.Presence of PCR products was assessed using 0.5% agarose (Sigma-Aldrich, USA, A9539) gel electrophoresis with SYBR Safe DNA Gel Stain (Invitrogen, USA, S33102), and 1 kb DNA ladders (New England Biolabs, N3232S). Amplicons were subsequently purified using the Genomic DNA Clean & Concentrator®-10 (D4011). Amplicon quantification and size assessment were performed using TapeStation 2200 (Agilent Technologies, G2965A) and the D5000 ScreenTape System (Agilent Technologies, 5067-5588, 5067-5589).

### PCR Barcoding

Barcoding PCR was performed using the PCR Barcoding Expansion 1-12 kit (Oxford Nanopore Technologies, EXP-PBC001). Products from the initial PCR were adjusted to a final volume of 48 μL using nuclease-free water. Only products with sufficiently high concentrations were adjusted to a final concentration of 4.17 nM. Each 100 μL reaction contained 48 μL of PCR product, 2 μL of 10 μM PCR barcode, and 50 μL of PCR mix consisting of 20 μL of 5X Phusion HF Buffer, 2 μL of 10 mM dNTP mix, 1 μL of Phusion Hot Start II High-Fidelity DNA Polymerase, and 27 μL of nuclease-free water. PCR conditions were initial denaturation at 98°C for 30 seconds, followed by 15 cycles of 98°C for 10 seconds, 62°C for 15 seconds, 72°C for 2 minutes, and final extension at 72°C for 10 minutes. Amplicons were purified using the Genomic DNA Clean & Concentrator®-10 (D4011) and quantified using the broad-range double-stranded DNA assay kit (Invitrogen, USA, Q32853) on the Qubit 2.0 Fluorometer (Invitrogen, USA, Q32866) and TapeStation 2200.

### Library preparation and sequencing

Barcoded PCR products were pooled in 47 μL of nuclease-free water at equal ratios (totalling 1 μg) and library preparation was carried out using the Ligation Sequencing Kit 1D (Oxford Nanopore Technologies, SQK-LSK109) following manufacturer’s protocols. Samples were quantified by Qubit 2.0 and TapeStation 2200. R9.4.1 flow cells (Oxford Nanopore Technologies, FLO-MIN106.001) were primed and quality controlled following manufacturer’s instructions prior to sample loading. Sequencing was performed using MinKNOW v22.03.5 with 36 hours runtime. Fast basecalling and barcoding was enabled and reads with Qscore < 8 were filtered out. Reads longer than 500 bp were retained for downstream analysis.

### Bioinformatics pipeline

Sequencing reads were adapter trimmed using Porechop (v0.2.4), with a min_split_read_size of 400 bp. Adapter-trimmed reads with lengths greater than 500 bp were retained for further analysis.

Taxonomic assignment of sequencing reads was carried out using Kraken2 (v.2.1.2), a classification system based on exact *k*-mer matches^28^. A custom Kraken2 database was built using genome assemblies at all levels for fungi obtained from RefSeq release 214^29^. The genomes of *C. dermatis* (ATCC 204094), *F. keratoplasticum* (ATCC 36031), and *T. interdigitale* (ATCC 9533), present in the Mycobiome Genomic DNA Mix but absent from RefSeq, were added to the database to evaluate the capabilities of Kraken’s species classification. Sankey diagrams were created using Pavian^30^. Bracken (v2.8), a companion program to Kraken2^31^, was used to estimate the abundance of each species and evaluate biodiversity. A Bracken database file was first generated using the Kraken2 database with *k*-mer length set to 35 and read length set to 500 bp. Sample data were passed through Bracken with the read length set to 500 bp, level for abundance estimation set to species, and threshold set to 10 reads. Bracken reports generated were then used to calculate α-diversity (Berger Parker’s, Fisher’s, Simpson’s, and Shannon’s index) and β-diversity (Bray-Curtis dissimilarity) using the KrakenTools python scripts^32^.

### Microbiology testing

All Human AD/PD brains and healthy control brains (Table 1) were swabbed in triplicate. Swabbing was performed with a sterile disposable loop into a plate of Sabouraud Dextrose Agar (SDA) containing 16 mg/L of Penicillin and Streptomycin. The same procedure was repeated with SDA containing chloramphenicol at concentration of 100 mg/L. All plates were sealed with parafilm and incubated at 37°C for 4 weeks.

### 16S ribosomal RNA PCR

Primers targeting the bacterial 16S ribosomal operon from position 357 to 926 (∽569bp) were used (Table 2 and Figure 2). PCR settings are shown in Table 2. PCR products were purified using the Genomic DNA Clean & Concentrator®-10 (D4011) and sent for Sanger DNA sequencing (GENEWIZ UK Ltd).

**Figure 2:**
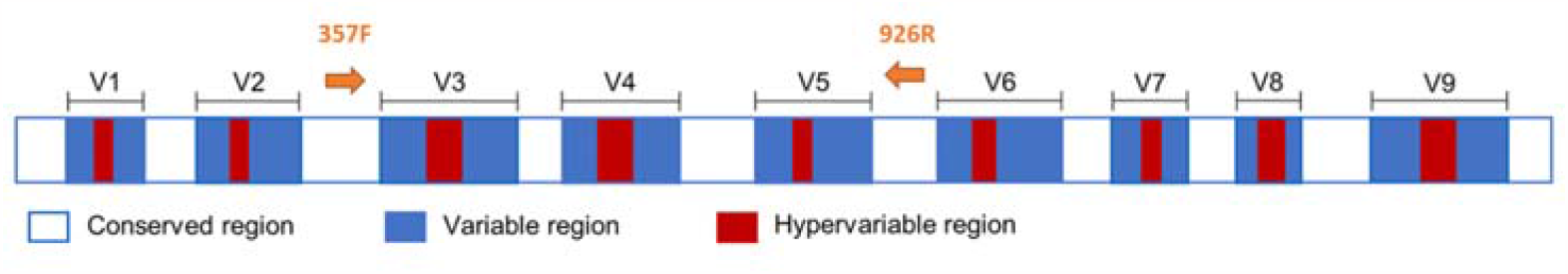
Conserved, variable, and hypervariable regions within the 16S ribosomal RNA. Forward primer 357F and Reverse primer 926R used in this study are represented in orange. Illustration adapted from Abellan-Schneyder et al.^33^

### Molecular analysis of bacteria

The 16S sRNA sequences generated during this study were aligned using ClustalW in Mega 11 by comparing it with seven bacterial reference sequences retrieved from GenBank: *Escherichia coli* (MN900682.1); *Pseudomonas aeruginosa* (FJ972538.1); *Pseudomonas* spp. (Strain AL208, MG819591.1); *Klebsiella pneumoniae* (AH009269.2); *Citrobacter freundii* (KC344791.1); *Staphylococcus aureus* (L37597.1); *Thermoplasma acidophilum* (M38637.1).

A phylogenetic tree based on eight nucleotide sequences was generated using a Maximum Likelihood method (1000 bootstrap, Kimura 2-parameter model^34–36^). *T. acidophilum* was used as an outgroup (Figure 3). Estimates of evolutionary divergence between the eight nucleotide sequences (Supplementary file S1) were calculated with MEGA 11 using a gamma distribution (shape parameter =1) with the following outgroup: *T. acidophilum*^35,37^.

**Figure 3:**
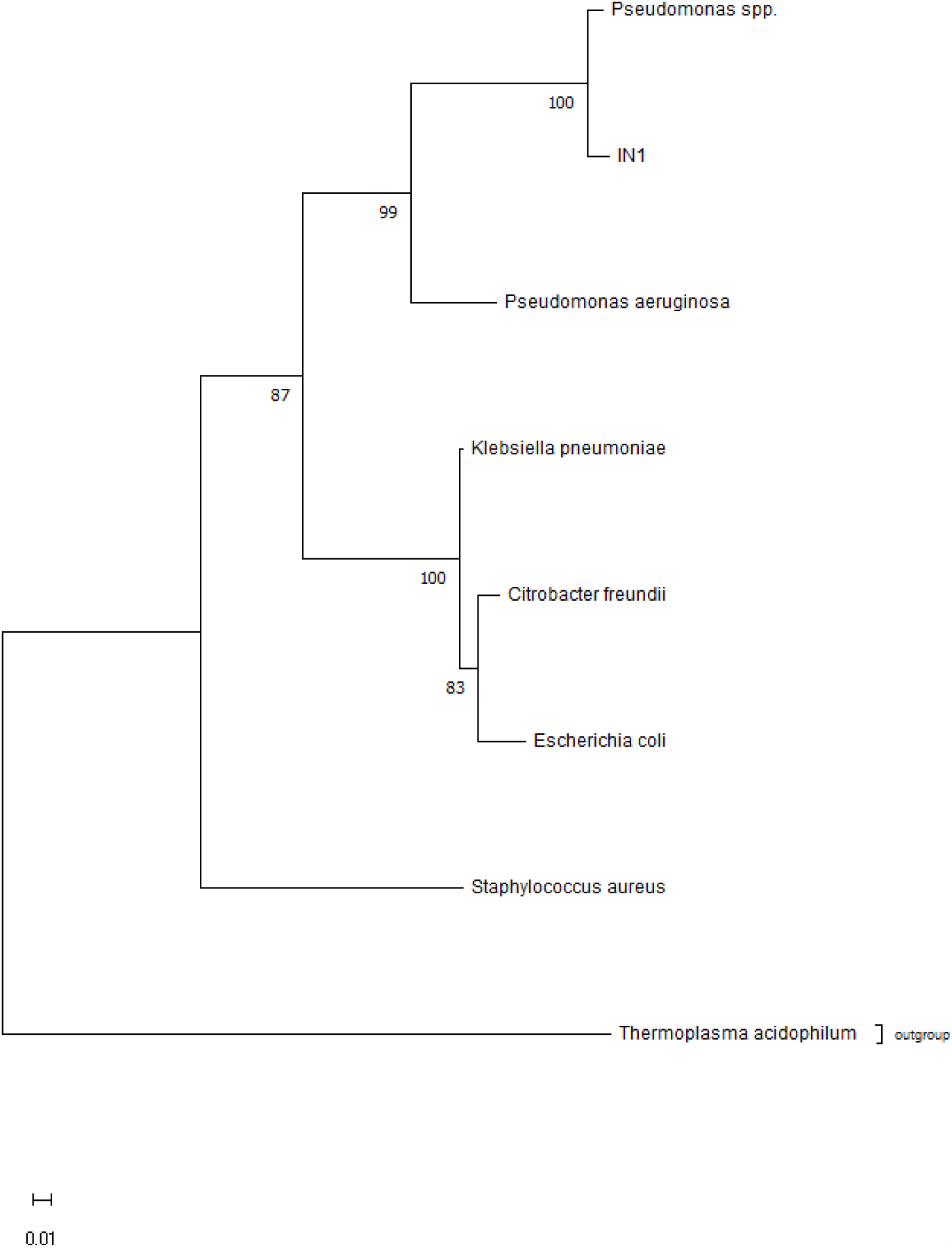
Evolutionary estimation analysis by the maximum likelihood method. The evolutionary history was inferred by using the maximum likelihood method and Kimura 2-parameter model [1]. The tree with the highest log likelihood (−8882.86) is shown. The percentage of trees in which the associated taxa clustered together is shown below the branches. Initial tree(s) for the heuristic search were obtained automatically by applying Neighbor-Join and BioNJ algorithms to a matrix of pairwise distances estimated using the Maximum Composite Likelihood (MCL) approach, and then selecting the topology with superior log likelihood value. The tree is drawn to scale, with branch lengths measured in the number of substitutions per site. This analysis involved eight nucleotide sequences. Codon positions included were 1st+2nd+3rd+Noncoding, generating a total of 2,998 positions in the final dataset. Evolutionary analyses were conducted in MEGA11 with 1000 bootstrap replications^34,35^.

## Results

### Quantitative polymerase chain reaction (qPCR)

qPCR was performed on DNA extracted from mice brain aliquots spiked with 10-fold serial dilutions of fungal spore mock community containing *A. fumigatus, C. auris*, and *Cryptococcus spp*. The limit of detection (LOD) was found to range between 100 and 1000 spores. Differences in LOD between assays may be dependent on qPCR efficiency which differs between assays as well as between sample replicates. For the 100 spore LOD qPCR assay, the following average Cq values apply: 37.28 (10,000 spores), 41.66 (1000 spores), and 43.12 (100 spores). There was no qPCR amplification for spiked brains with 10 spores, 1 spore, and 0 spores. We observed no amplification for the no template control, nuclease free water, and PBS. Upon analysing all human brain samples and CSF (Table 1) with the FungiQuant assay we detected no fungal DNA. The efficiency of the qPCR assays was 81.98%, 87.23%, and 90.989% with R^2^ values of 0.98, 0.989, and 0.92 respectively.

### Reverse transcription (RT)

RNA extracted from human brain samples (Table 1) was transcribed into cDNA and a qPCR assay was performed using the 18S primers mentioned above. No fungal cDNA was detected in any of the brain samples (Table 1).

### Microbiological testing

All human brain samples (Table 1) were swabbed and inoculated onto SDA plates containing 16 mg/L of penicillin and streptomycin and 100 mg/L of chloramphenicol. No fungal growth was observed after four weeks on any plate. However, we observed bacterial growth for sample IN1, which is a microbiological swab from sample AD2_1 (Table 1) belonging to an AD patient. We then confirmed our finding by PCR using 16S rRNA primers (Table 2) and performing species identification. Submitting the 16S sRNA partial sequence of sample IN1 into BLAST^38^ identified the bacteria as *Pseudomonas spp*., with an E value of 0 and percent identity of 97.82% (program version: BLASTN 2.14.1+).

Phylogenetic analysis showed sample IN1 clustering on the same branch as *P. aeruginosa* with a high bootstrap support of 99 (Figure 3). This finding is further supported by the estimation of evolutionary divergence between *Pseudomonas spp*. and IN1, identified as 0.0164 (which is the lowest P-Distance of all 16S sRNA sequences used in this analysis), whereas the distance between IN1 and the outgroup *T. acidophilum* was 0.3856 (supplementary Table 1). We decided to remove all 16S rRNA sequences generated during this study except for sample IN1 as Sanger sequencing was terminated early or was non-specific for all sequences except for sample IN1. One possible explanation for earlier termination or non-specific results during Sanger sequencing might be the presence of mixed colonies of bacteria in the human brain tissues, whereas IN1 was isolated on a SAD plate followed by DNA extraction and PCR.

### Kraken2 species classification

The Mycobiome Genomic DNA Mix was sequenced on MinION and analysed using Kraken2 to assess pipeline sensitivity. A total of 66,141 reads were classified and eight out of ten species were detected, with the exception of *A. fumigatus* and *P. chrysogenum*, potentially due to low abundance or high sequence similarity to other species (Figure 4). All ten genera present in the mock community were identified, albeit at unequal proportions (Figure 4).

**Figure 4:**
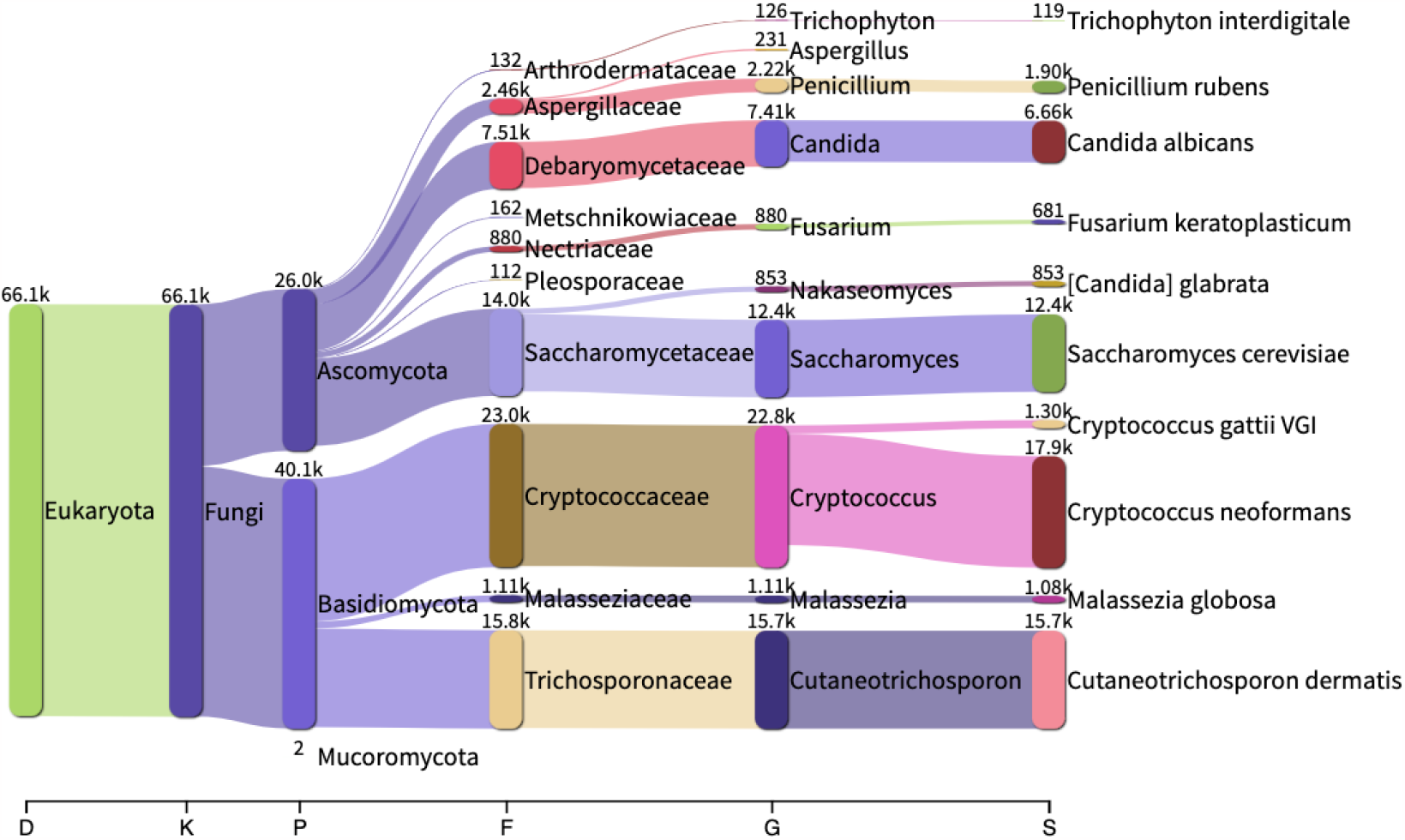
Sankey diagrams depicting Kraken2 classification results and the top ten most abundant species in terms of read count obtained from the Mycobiome Genomic DNA Mix at 10 genome copy equivalence.

Human brain and CSF samples, mock communities, and ‘DNA-free’ negative controls (BSA and DNA extraction kit control) were sequenced on MinION and analysed using Kraken2 to determine which species, if any, are present in AD/PD and control brains and CSF and assess the level of noise or contamination originating from ‘DNA-free’ controls. The number of total classified reads for each sample are shown in Table 3. Species reads obtained from BSA in each sequencing run (BSA 1 for samples 0-12, BSA 2 for samples 13-19) were subtracted from those obtained from each sample and these processed read counts were used to visualise the proportions of reads obtained from the most abundant species identified by Kraken2 (Figure 5). Amongst the top ten species identified across all samples, *Alternaria ethzedia, Alternaria hordeiaustralica*, and *Alternaria metachromatica* were consistently present in all human samples (Figure 5A), absent in mock community samples (Figure 5B), and present at lower levels in ‘DNA-free’ controls (Figure 5C). Their almost ubiquitous detection could indicate these species as laboratory contaminants, however the possibility of their presence in some samples cannot be eliminated as samples 5 (AD brain), 6 (control brain with extensive amyloid angiopathy), and 8 (AD brain) had high numbers of total classified reads (21,373, 10,207, and 21,741 reads respectively) and *Alternaria* reads constitute significant proportions of these reads, ranging from approximately 14% to over 50% (Figure 5A). Samples 5, 6, and 8 also contain reads from *Colletotrichum graminicola*, at approximately 6%, 16%, and 2% of all classified reads. *Filobasidium floriforme* appears to be present at relatively high proportions in samples 1, 5, and 8, constituting approximately 47%, 57%, and 13% of all classified reads respectively (Figure 5A). The species makeup of sample 9, CSF obtained from the same patient as sample 8, differs from its brain counterpart, containing higher proportions of reads from *Malassezia spp*. Samples 12 to 16 contained high proportions of reads from *C. auris, Cryptococcus spp*., and *A. fumigatus* (with the exception of sample 12), likely due to contamination during DNA handling or problems during demultiplexing, as these samples were sequenced in the same run with the mock community containing the above species, and the species makeup of these samples is similar to that of BSA 2 (sample 17), which was also sequenced in the same run (Figure 5).

**Table 3:** Total number of Kraken2 classified reads. Sequencing runs are separated by a bolded black line between samples 11 and 12.

| Code | Sample | Sample type | Condition | Total classified reads |
| --- | --- | --- | --- | --- |
| 0 | C2.1 | Brain | Control | 4,674 |
| 1 | C2.2 | Brain | Control | 8,452 |
| 2 | C2.3 | Brain | Control | 2,558 |
| 3 | C1 | Brain | Control | 4,177 |
| 4 | AD1_1 | Brain | Disease | 675 |
| 5 | AD1_2 | Brain | Disease | 21,373 |
| 6 | C3_1 | Brain | Control | 10,207 |
| 7 | C3_2 | CSF | Control | 855 |
| 8 | AD2_1 | Brain | Disease | 21,741 |
| 9 | AD2_2 | CSF | Disease | 3,396 |
| 10 | BSA 1 | 'DNA free' | Standard | 1,226 |
| 11 | Mycobiome Genomic DNA Mix | Mock community | Standard | 66,141 |
| 12 | PD1 | Brain | Disease | 940 |
| 13 | AD3 | Brain | Disease | 1,504 |
| 14 | C4 | Brain | Control | 1,084 |
| 15 | PD2 | Brain | Disease | 1,548 |
| 16 | AD4 | Brain | Disease | 1,029 |
| 17 | BSA 2 | 'DNA free' | Standard | 135 |
| 18 | DNA extraction kit control | 'DNA free' | Standard | 631 |
| 19 | *C. auris*, *A. fumigatus*,<br>*Cryptococcus* spp. | Mock community | Standard | 88,386 |

**Figure 5:**
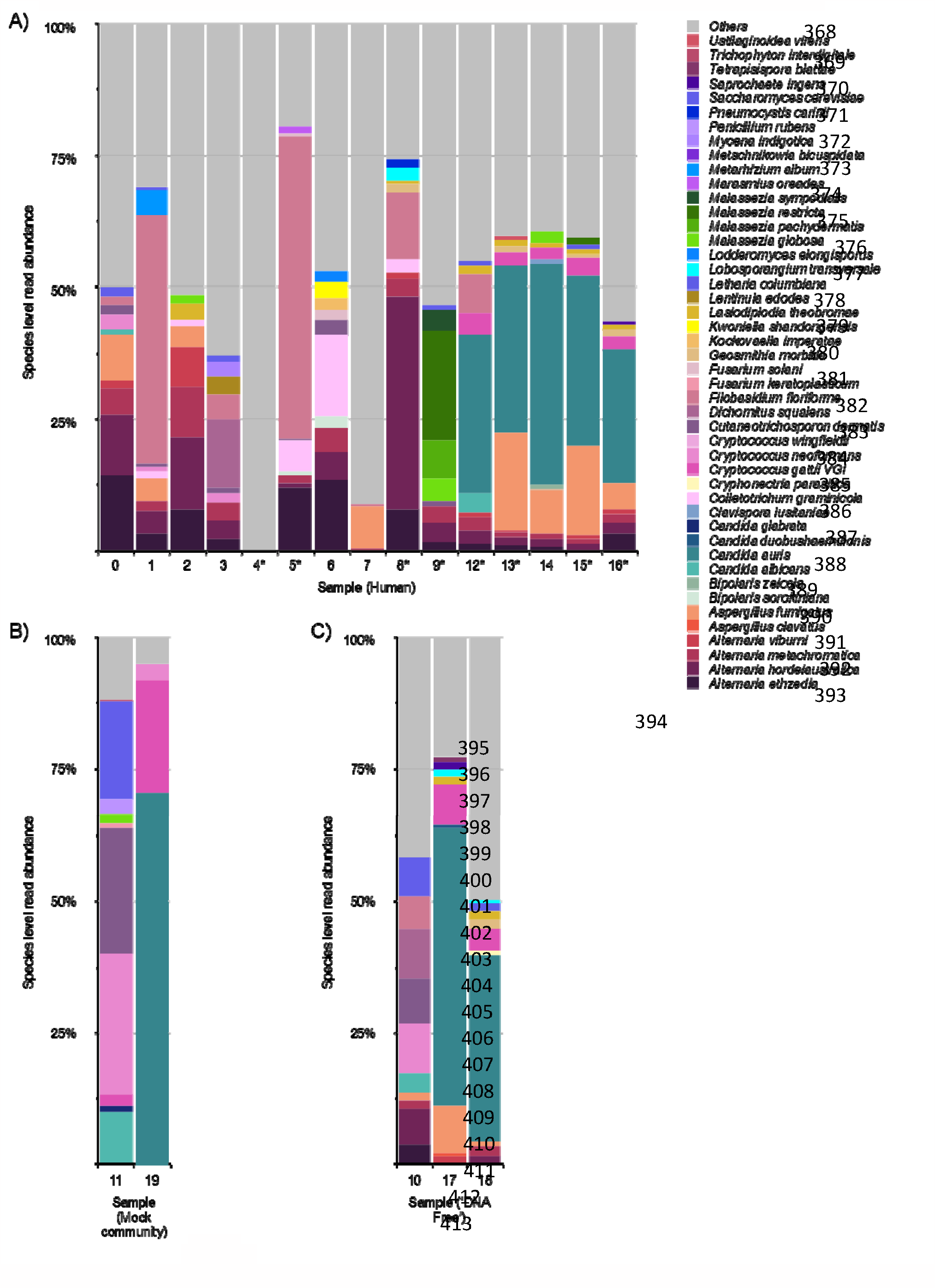
Species level read abundance in A) human samples, B) mock communities, and C) ‘DNA free’ controls based on Kraken2. Proportions calculated based on processed read counts over the total number of classified reads in each sample. Asterisks mark samples from AD/PD patients.

### α-diversity and β-diversity

No significant differences in within sample species diversity were found across control, disease, mock community, and ‘DNA free’ sample groups using different α-diversity measures such as Berger β-diversity based on the Bray-Curtis dissimilarity matrix was visualised using principal coordinate analysis (PCoA) (Figure 7). We found significant differences between control brains, disease brains, control CSF, disease CSF, ‘DNA free’ controls, and mock community samples (*p*=0.029, pseudo-F=1.547778, PERMANOVA), with principal coordinates 1 and 2 accounting for 32.39% and 18.03% of variance respectively (Figure 7). Control brains and disease brains form distinct clusters with the exception of some samples clustering with the opposite group. Mock community samples cluster closely and there is greater variation in ‘DNA free’ controls potentially due to different levels of contamination. Control and disease CSF samples are also clearly separated, however there is overlap with the control brain cluster as well as a disease brain sample. Of note, there is significant divergence between samples 4 and 5, corresponding to the cerebellum and frontal cortex from the same AD patient, as well as samples 6 and 7, corresponding to the frontal cortex and CSF from a control patient.

**Figure 6:**
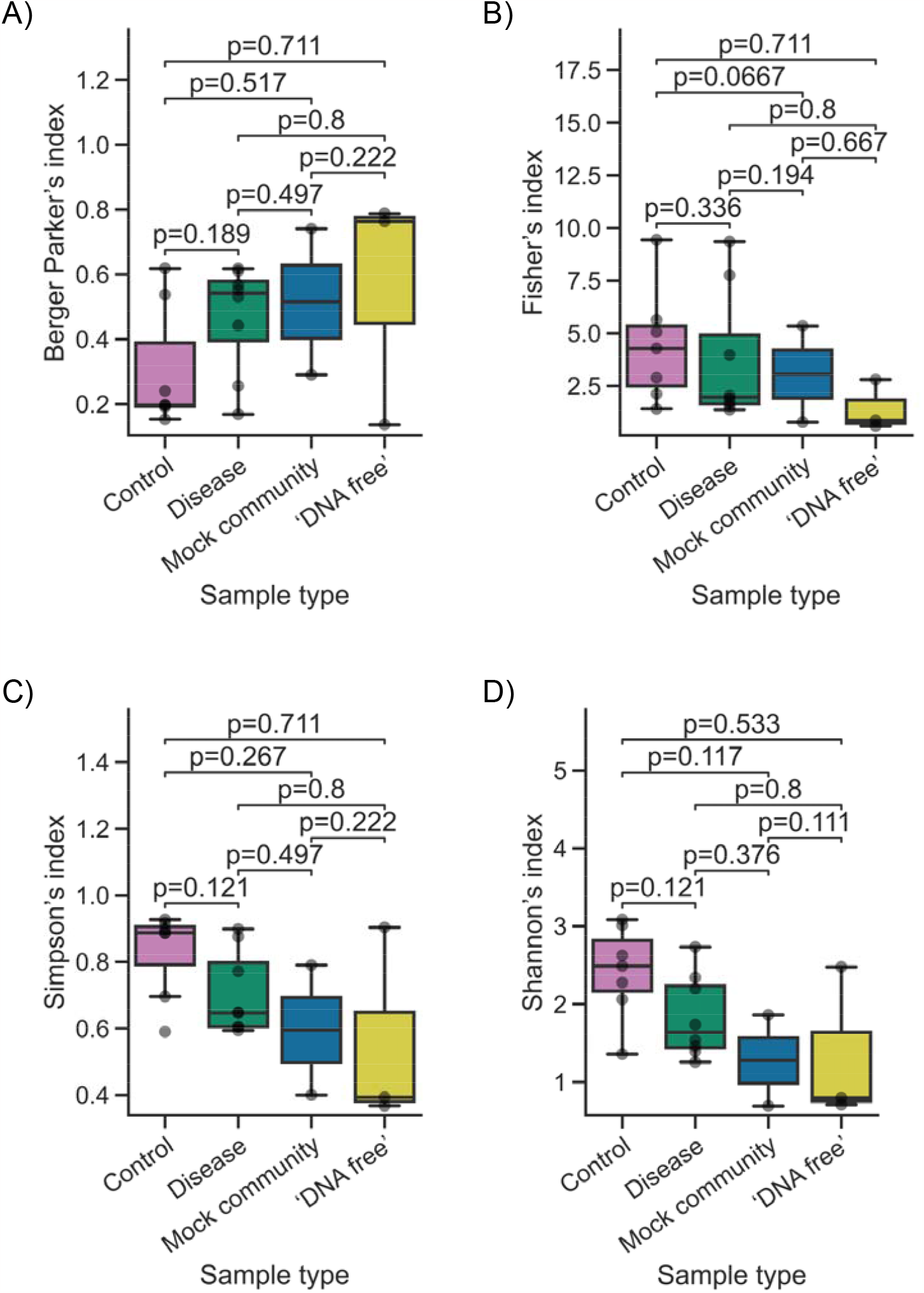
α-diversity among different sample groups or conditions measured using the Berger Parker’s index (A), Fisher’s index (B), Simpson’s index (C), and Shannon’s index (D). Whiskers extend to 1.5 times the interquartile range. *P*-values determined by two-sample Mann-Whitney U test comparing each sample group or condition.

**Figure 7:**
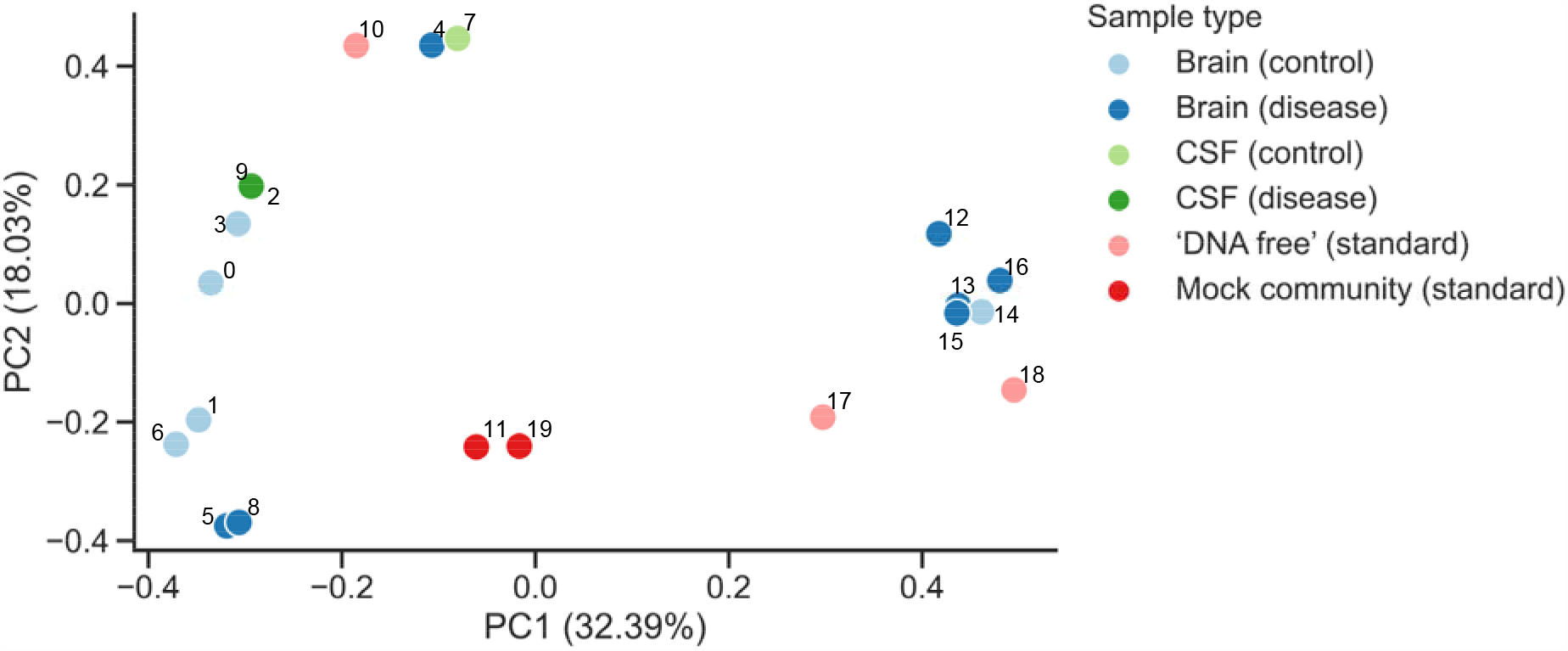
β-diversity PCoA plot based on a Bray-Curtis dissimilarity matrix generated by Bracken. Data points labelled according to sample numbers denoted in Table 3.

AD and PD are both neurodegenerative diseases associated with the abnormal aggregation of proteins in the CNS such as amyloid-β, tau and α-synuclein^39^. Despite the fact that both AD/PD have been studied for over a century, the exact causes of these neurodegenerative diseases are still far from being understood^40^. Several studies have been able to establish a link between bacterial or viral infection and AD/PD^13,14^, however, few have attempted to establish a connection between fungal infection and AD/PD. Studies conducted by Pisa and Alonso *et al*.^17–22^ have detected fungal DNA in AD/PD brains belonging to species in the *Alternaria, Botrytis, Candida, Cladosporium, Malassezia, Saccharomyces*, and *Cryptococcus* genera, with *Cryptococcus* being the most commonly found genus. *Alternaria, Candida, Cladosporium, Malassezia*, and *Cryptococcus* species have been found to be capable of infecting the CNS and are capable of causing fungal menigitis^16,42^. In fact, there have been reports of Cryptococcal meningitis misdiagnosed as AD due to its cause of dementia-like symptoms in patients, who demonstrated loss of memory and independence in performing certain activities as well as slight behavioural changes, with these cognitive deficits reversed following antifungal treatment^41,42^.

Here, we used standard PCR-based methods alongside nanopore sequencing on the MinION platform to profile the microbial content of healthy and AD/PD brains and CSF. First, we developed and validated a method allowing the detection of fungal DNA in mammalian brain tissue. Primers targeting the fungal ribosomal operon with an amplicon size of 3.5 kb were used, followed by nanopore sequencing, as well as qPCR and RT-qPCR (Figure 1 and Table 2). qPCR and RT-qPCR did not detect fungal DNA in any human brain/CSF samples. Secondly, we used MinION sequencing to detect and quantify fungal species present in brain tissue samples. We found evidence of fungal DNA in both healthy and AD/PD human brain samples attributed to *Alternaria spp*., *C. graminicola*, and *F. floriforme*. Our findings are in line with previous studies conducted by Alonso *et al*.^20,21^, where they also reported the presence of fungal DNA on healthy human brains as well AD/PD brains. This finding further supports the usage of third generation technologies such as nanopore sequencing as a very valuable tool for detecting low levels of fungal DNA in human brain samples over traditional methods such as (q)PCR.

Nanopore sequencing proves more sensitive to the detection of fungal DNA in brain samples even at low levels, whilst FungiQuant failed to detect any DNA; this may be explained by the variation of LOD (which varied between 100-1000 spores). Therefore, if the brain sample had less than 100 fungal genomes present the FungiQuant assay would not be able detect it. To our knowledge, this is a novel application of the MinION sequencer, allowing rapid characterisation of microbial content in CNS tissue samples.

Long-read sequencing provided by the MinION platform enabled improved taxonomic classification. This in turn provided species-level information despite the slightly higher error rate compared to short-read sequencing^43^. However, sequencing errors generated in barcode sequences may affect accurate demultiplexing, demonstrated by the contamination of human samples by mock community species (Figure 5). Additionally, though the *k*-mer based classification employed by Kraken2 is highly efficient, the increased error rate of long reads may result in misclassification of closely related species^43^. Kraken2 also relies on a well-curated taxonomy tree from the National Center for Biotechnology Information for classification, which may not contain nodes for newly discovered species^44^.

Highly abundant fungal species identified across multiple patient samples included *Alternaria spp*., *C. graminicola*, and *F. floriforme. Alternaria* is known to be able to infect the CNS and is also a part of the human microbiome^15,42,45^. *C. graminicola* has been reported as a human pathogen in rare cases, causing ocular infections^46,47^. *F. floriforme* itself has not been reported as a human pathogen, but closely related species such as *Filobasidium uniguttulatum* have been shown to cause meningitis^48^. Identification of these species could suggest their presence in patient brains as a part of the microbiome or as a result of infection, however links to neurodegenerative diseases remain unclear since these species were found in both control and disease patient samples and differences in abundances cannot be evaluated accurately due to different PCR or barcoding efficiencies. Our analysis of α-diversity did not reveal significant differences in species richness or evenness across different sample groups, which could be attributed to our relatively small sample size as well as high levels of contamination in one of the sequencing runs. β-diversity analysis, on the other hand, revealed differences between sample groups and within some patients in terms of the number of species present and their abundance, however, this can also be affected by contamination caused by DNA handling or demultiplexing. Furthermore, Bracken’s abundance estimation may not translate well into Nanopore sequencing data generated with the use of PCR and results from diversity analysis must be interpreted with caution. Future studies should endeavour to employ PCR-free sequencing methods for more accurate abundance estimations and biodiversity analysis.

Although the initial focus of this study was to investigate the presence of fungi in human brain tissue and CSF, an antibiotic resistant bacterium was also detected when assessing brain AD2_1 (Table 1). Assigned IN1, after Sanger sequencing we identified the resistant bacterium as a member of the *Pseudomonas* genus. It has been reported that *P. aeruginosa* may increase tau hyperphosphorylation, which is a hallmark of AD^49^. Hence, it is possible that IN1 could have contributed to the development of AD within this patient. However, further studies are needed to strengthen our confidence in this relationship. It is challenging to determine whether IN1 was the result of a brain infection or the consequence of physical contamination during initial dissection of the neurological tissue or during its handling in the laboratory.

To improve sensitivity and confidence of species identification in metabarcoding studies, multiple pipelines and databases may need to be used, and newer machine learning-based classification systems may also be considered to enable the identification of novel species^44^. Use of fungi-specific primers for PCR also reduces the need for depletion of background or host DNA and can increase detection of species with lower abundances^43^. This is however accompanied by the drawback of PCR artefacts and biases which affect the estimation of species abundance as the use of universal primers produces different amplification efficiencies in different species^43^. Varying copy numbers of the ribosomal operon across species may also affect the number of reads obtained from each species and skew abundance estimations^43^.

In conclusion, this study observed the presence of fungal DNA in healthy human brains, as well as AD/PD brains. The protocol developed to detect low levels of fungal DNA in CNS tissue will be of use for further exploring the microbial aetiology hypothesis for AD and PD.

## Data availability

All raw reads have been submitted to the European Nucleotide Archive under accession ERP150444.

## Code availability

All scripts can be found at https://github.com/mycologenomics/AD_PD_microbiome.

## Acknowledgements

This paper presents independent research funded by a Wellcome Trust Institutional Strategic Support Fund’s Springboard Fellowship Scheme (Imperial College London) awarded to JR (WPIA_PSN105). RL and MCF were supported by a grant from the Wellcome Trust (219551/Z/19/Z). Dr. Tom Williams and Professor Darius Armstrong-James (DIDE, Imperial college) acknowledged for the donation of most the mice brain samples analysed in this study. In addition, we also would like to acknowledge Dr. Norman van Rhijn and Professor Mike Bromley (Manchester Fungal Infection Group) for the donation of mice brains used during this project. JR has previously received an honoraria from Gilead Sciences. MCF is funded by the CIFAR Fungal Kingdoms program.

## Author Contributions

R.L, M.C.F and J.R conceived and designed the study. R.L and I.U.W carried out the experimental work. R.L, I.U.W and J.R carried out the experimental validation and analysis. R.L, I.U.W, M.C.F and J.R wrote the paper.

## Supplementary information

Supplementary File S1 and Supplementary Table 1.

## References

1. Ransohoff, R. M. How neuroinflammation contributes to neurodegeneration. Science 353, 777–783 (2016).

2. Scheltens, P. et al. Alzheimer’s disease. Lancet 397, 1577–1590 (2021).

3. Alzheimer’s disease | Alzheimer’s Society. https://www.alzheimers.org.uk/about-dementia/types-dementia/alzheimers-disease (2023).

4. Keener, A. M. & Bordelon, Y. M. Parkinsonism. Semin Neurol 36, 330–334 (2016).

5. Walker, Z., Possin, K. L., Boeve, B. F. & Aarsland, D. Lewy body dementias. Lancet 386, 1683–1697 (2015).

6. alzheimer_europe_dementia_in_europe_yearbook_2019.pdf. https://www.alzheimereurope.org/sites/default/files/alzheimer_europe_dementia_in_europe_yearbook_2019.pdf.

7. Lindestam Arlehamn, C. S., Garretti, F., Sulzer, D. & Sette, A. Roles for the adaptive immune system in Parkinson’s and Alzheimer’s diseases. Curr Opin Immunol 59, 115–120 (2019).

8. Tahami Monfared, A. A., Byrnes, M. J., White, L. A. & Zhang, Q. Alzheimer’s Disease: Epidemiology and Clinical Progression. Neurol Ther 11, 553–569 (2022).

9. Long, J. M. & Holtzman, D. M. Alzheimer Disease: An Update on Pathobiology and Treatment Strategies. Cell 179, 312–339 (2019).

10. Kurz, C., Walker, L., Rauchmann, B.-S. & Perneczky, R. Dysfunction of the blood-brain barrier in Alzheimer’s disease: Evidence from human studies. Neuropathol Appl Neurobiol 48, e12782 (2022).

11. Tansey, M. G. et al. Inflammation and immune dysfunction in Parkinson disease. Nat Rev Immunol 22, 657–673 (2022).

12. Shen, C.-H. et al. Association Between Tuberculosis and Parkinson Disease: A Nationwide, Population-Based Cohort Study. Medicine (Baltimore) 95, e2883 (2016).

13. Panza, F., Lozupone, M., Solfrizzi, V., Watling, M. & Imbimbo, B. P. Time to test antibacterial therapy in Alzheimer’s disease. Brain 142, 2905–2929 (2019).

14. Bu, X.-L. et al. A study on the association between infectious burden and Alzheimer’s disease. European Journal of Neurology 22, 1519–1525 (2015).

15. Phuna, Z. X. & Madhavan, P. A closer look at the mycobiome in Alzheimer’s disease: Fungal species, pathogenesis and transmission. Eur J Neurosci 55, 1291–1321 (2022).

16. Alonso, R., Pisa, D., Rábano, A., Rodal, I. & Carrasco, L. Cerebrospinal Fluid from Alzheimer’s Disease Patients Contains Fungal Proteins and DNA. J Alzheimers Dis 47, 873–876 (2015).

17. Pisa, D., Alonso, R., Juarranz, A., Rábano, A. & Carrasco, L. Direct visualization of fungal infection in brains from patients with Alzheimer’s disease. J Alzheimers Dis 43, 613–624 (2015).

18. Pisa, D., Alonso, R., Rábano, A., Rodal, I. & Carrasco, L. Different Brain Regions are Infected with Fungi in Alzheimer’s Disease. Sci Rep 5, 15015 (2015).

19. Pisa, D., Alonso, R., Rábano, A., Horst, M. N. & Carrasco, L. Fungal Enolase, β-Tubulin, and Chitin Are Detected in Brain Tissue from Alzheimer’s Disease Patients. Front Microbiol 7, 1772 (2016).

20. Alonso, R., Pisa, D., Aguado, B. & Carrasco, L. Identification of Fungal Species in Brain Tissue from Alzheimer’s Disease by Next-Generation Sequencing. J Alzheimers Dis 58, 55–67 (2017).

21. Alonso, R., Pisa, D., Fernández-Fernández, A. M. & Carrasco, L. Infection of Fungi and Bacteria in Brain Tissue From Elderly Persons and Patients With Alzheimer’s Disease. Front Aging Neurosci 10, 159 (2018).

22. Pisa, D., Alonso, R. & Carrasco, L. Parkinson’s Disease: A Comprehensive Analysis of Fungi and Bacteria in Brain Tissue. Int J Biol Sci 16, 1135–1152 (2020).

23. Link, C. D. Is There a Brain Microbiome? Neurosci Insights 16, 26331055211018709 (2021).

24. Verheggen, I. C. M. et al. Increase in blood-brain barrier leakage in healthy, older adults. Geroscience 42, 1183–1193 (2020).

25. Liu, C. M. et al. FungiQuant: a broad-coverage fungal quantitative real-time PCR assay. BMC Microbiol 12, 255 (2012).

26. Charalampous, T. et al. Nanopore metagenomics enables rapid clinical diagnosis of bacterial lower respiratory infection. Nat Biotechnol 37, 783–792 (2019).

27. D’Andreano, S., Cuscó, A. & Francino, O. Rapid and real-time identification of fungi up to species level with long amplicon nanopore sequencing from clinical samples. Biol Methods Protoc 6, bpaa026 (2021).

28. Wood, D. E., Lu, J. & Langmead, B. Improved metagenomic analysis with Kraken 2. Genome Biol 20, 257 (2019).

29. O’Leary, N. A. et al. Reference sequence (RefSeq) database at NCBI: current status, taxonomic expansion, and functional annotation. Nucleic Acids Research 44, D733–D745 (2016).

30. Breitwieser, F. P. & Salzberg, S. L. Pavian: interactive analysis of metagenomics data for microbiome studies and pathogen identification. Bioinformatics 36, 1303–1304 (2020).

31. Lu, J., Breitwieser, F. P., Thielen, P. & Salzberg, S. L. Bracken: Estimating species abundance in metagenomics data. PeerJ Computer Science 2017, e104 (2017).

32. Lu, J. et al. Metagenome analysis using the Kraken software suite. Nat Protoc 17, 2815–2839 (2022).

33. Abellan-Schneyder, I. et al. Primer, Pipelines, Parameters: Issues in 16S rRNA Gene Sequencing. mSphere 6, 10.1128/msphere.01202-20 (2021).

34. Kimura, M. A simple method for estimating evolutionary rates of base substitutions through comparative studies of nucleotide sequences. J Mol Evol 16, 111–120 (1980).

35. Tamura, K., Stecher, G. & Kumar, S. MEGA11: Molecular Evolutionary Genetics Analysis Version 11. Molecular Biology and Evolution 38, 3022–3027 (2021).

36. Felsenstein, J. CONFIDENCE LIMITS ON PHYLOGENIES: AN APPROACH USING THE BOOTSTRAP. Evolution 39, 783–791 (1985).

37. Tamura, K., Nei, M. & Kumar, S. Prospects for inferring very large phylogenies by using the neighbor-joining method. Proc Natl Acad Sci U S A 101, 11030–11035 (2004).

38. Pseudomonas sp. strain AL208 16S ribosomal RNA gene, partial sequence - Nucleotide - NCBI.https://www.ncbi.nlm.nih.gov/nucleotide/MG819591.1?report=genbank&log$=nuclalign&blast_rank=1&RID=D23XPBTB016.

39. Mucke, L. Neuroscience: Alzheimer’s disease. Nature 461, 895–897 (2009).

40. Zhang, W., Xiao, D., Mao, Q. & Xia, H. Role of neuroinflammation in neurodegeneration development. Signal Transduct Target Ther 8, 267 (2023).

41. Ala, T. A., Doss, R. C. & Sullivan, C. J. Reversible dementia: a case of cryptococcal meningitis masquerading as Alzheimer’s disease. J Alzheimers Dis 6, 503–508 (2004).

42. Hoffmann, M., Muniz, J., Carroll, E. & De Villasante, J. Cryptococcal meningitis misdiagnosed as Alzheimer’s disease: complete neurological and cognitive recovery with treatment. J Alzheimers Dis 16, 517–520 (2009).

43. Ciuffreda, L., Rodríguez-Pérez, H. & Flores, C. Nanopore sequencing and its application to the study of microbial communities. Comput Struct Biotechnol J 19, 1497–1511 (2021).

44. Q, L., Pw, B. Y L. B Z. & L W. DeepMicrobes: taxonomic classification for metagenomics with deep learning. NAR genomics and bioinformatics 2, (2020).

45. Seed, P. C. The human mycobiome. Cold Spring Harb Perspect Med 5, a019810 (2014).

46. Ritterband, D. C., Shah, M. & Seedor, J. A. Colletotrichum graminicola: a new corneal pathogen. Cornea 16, 362–364 (1997).

47. Yegneswaran, P. P., Pai, V., Bairy, I. & Bhandary, S. Colletotrichum graminicola keratitis: First case report from India. Indian J Ophthalmol 58, 415–417 (2010).

48. Pan, W. et al. Meningitis caused by Filobasidium uniguttulatum: case report and overview of the literature. Mycoses 55, 105–109 (2012).

49. Ochoa, C. D., Alexeyev, M., Pastukh, V., Balczon, R. & Stevens, T. Pseudomonas aeruginosa Exotoxin Y Is a Promiscuous Cyclase That Increases Endothelial Tau Phosphorylation and Permeability. J Biol Chem 287, 25407–25418 (2012).

