## Supplementary File S1 for "Determining the microbial species content in tissue from Alzheimer’s and Parkinson’s Disease patients"

**Supplementary File S1. DNA extraction method and validation qPCR (LOD).**

5 different DNA extraction methods were tested during this study, in order to understand which method would be the most efficient at extracting genomic DNA from mammalian brain tissue. A mock community of 3 fungal species (*Aspergillus fumigatus*, *Candida albicans* and *Cryptococcus* spp) was used as positive control and total of 600 spores were spiked into mice brains. Mice brains were incubated at 37°C for 17 hours in a rotatory incubator at 300rpm. Brains were centrifuged at maximum speed and the supernatant was discarded. DNA extractions were performed following the 5 protocols mentioned below:

1. DNeasy Blood & Tissue Kit with no bead beating

2. DNeasy Blood & Tissue Kit with beading (PowerBead Pro Tubes, Cat. No 19301 before addition of Proteinase K (Step 2)

3. DNeasy Blood & Tissue Kit with beading (PowerBead Pro Tubes, Cat. No 19301) after addition of Proteinase K (Step 3)

4. DNeasy PowerSoil Pro Kits (Qiagen, Germany, 47014)

5. ZymoBIOMICS DNA Miniprep Kit (D4300)

Average Cq values from FungiQuant (qPCR assay) were used to determine which method performed better in relation to the others. The DNA extraction method that amplified earlier on the qPCR was selected. DNeasy PowerSoil Pro Kits performed better than ZymoBIOMICS DNA Miniprep Kit (method 5) and DNeasy Blood & Tissue Kit (method 1) with Cq values of: 28.84; 29.14 and 34.59 respectively. Method 2 and 3 did not perform as well as the others and were removed from the analysis.

**Supplementary Table 1. Estimates of Evolutionary Divergence between Sequences.**

|  | Pseudomonas_aeruginosa | Thermoplasma_acidophilum_{outgroup} | Citrobacter_freundii | Klebsiella_pneumoniae | Staphylococcus_aureus | Pseudomonas_sp. | Escherichia_coli | IN1 |
| --- | --- | --- | --- | --- | --- | --- | --- | --- |
| Pseudomonas_aeruginosa |  |  |  |  |  |  |  |  |
| Thermoplasma_acidophilum_{outgroup} | 0.3856 |  |  |  |  |  |  |  |
| Citrobacter_freundii | 0.1611 | 0.3865 |  |  |  |  |  |  |
| Klebsiella_pneumoniae | 0.1708 | 0.3934 | 0.0259 |  |  |  |  |  |
| Staphylococcus_aureus | 0.2376 | 0.4007 | 0.2285 | 0.2315 |  |  |  |  |
| Pseudomonas_sp. | 0.1288 | 0.4045 | 0.1977 | 0.2181 | 0.2445 |  |  |  |
| Escherichia_coli | 0.1700 | 0.3787 | 0.0384 | 0.0414 | 0.2330 | 0.2160 |  |  |
| IN1 | 0.0764 | 0.3757 | 0.1585 | 0.1364 | 0.2113 | 0.0164 | 0.1621 |  |

The number of base differences per site from between sequences are shown. The rate variation among sites was modeled with a gamma distribution (shape parameter = 1). This analysis involved 8 nucleotide sequences. Codon positions included were 1st+2nd+3rd+Noncoding. All ambiguous positions were removed for each sequence pair (pairwise deletion option). There were a total of 2998 positions in the final dataset. Evolutionary analyses were conducted in MEGA11 [1]

**Supplementary References**

1. Tamura K., Stecher G., and Kumar S. (2021). MEGA 11: Molecular Evolutionary Genetics Analysis Version 11. Molecular Biology and Evolution https://doi.org/10.1093/molbev/msab120.
